# Neutral pH Plasma-Activated Water eliminates ESKAPE pathogens via reactive nitrogen species with dual potential for surface sanitation and wound cleansing

**DOI:** 10.1101/2024.10.06.616848

**Authors:** Debapriya Mukherjee, Pallab Ghosh, Atish Roy Chowdhury, Nishanth Vishwa, Kavita Verma, Lakshminarayana Rao, Dipshikha Chakravortty

## Abstract

Plasma-activated water (PAW) is an emerging antimicrobial strategy, though its efficacy has traditionally depended on acidic pH to generate reactive species. To improve clinical and practical applicability, we developed a neutral pH formulation using buffered water—hs-PAbW—and evaluated its antimicrobial activity and mechanism of action against ESKAPE pathogens. hs-PAbW significantly reduced bacterial viability during the exponential growth phase. While reactive oxygen species (ROS) alone were insufficient for bacterial killing, reactive nitrogen species (RNS), particularly peroxynitrite, mediated strong microbicidal effects via nitrotyrosine formation. These effects occurred independently of acidic conditions, demonstrating that antimicrobial activity is retained at neutral pH. Alongside, hs-PAbW effectively reduced bacterial burden on metal surfaces and exhibited superior killing compared to high doses of standard antibiotics, including ceftazidime, ciprofloxacin, and meropenem *in vitro*. A non-cytotoxic dose of hs-PAbW also reduced bacterial load in infected wounds and accelerated wound closure *in vivo*. These findings highlight the dual functionality of hs-PAbW in surface sanitation and wound care, and support its potential as a sustainable, broad-spectrum antimicrobial agent for clinical use.

## Introduction

Antimicrobial resistance (AMR) is recognized by the World Health Organization as a critical global health threat, with over two million antibiotic-resistant infections and 23,000 related deaths annually in the US alone^1, 2^. Globally, bacterial AMR caused 1.27 million deaths in 2019^3^. *Acinetobacter*, *Pseudomonas,* and non-typhoidal *Salmonella* are a few of the bacterial pathogens which are predominantly associated with AMR-related deaths ^1, 3, 4^. This is why developing a novel, long-lasting, and sustainable solution for bacterial infections remains a significant necessity.

Plasma-activated water (PAW) is an emerging technology that has gained attention for its potential as a sustainable solution to various medical and food preservation applications ^5–7^. PAW is produced by exposing water to cold plasma, an ionized gas containing charged particles like ions, electrons, and neutral atoms ^8^. The ionized particles in the plasma interact with the water molecules generating reactive oxygen species (ROS) and reactive nitrogen species (RNS)^9^. These ROS and RNS generated have varying lifetimes, ranging from a few nanoseconds to a few years ^10, 11^. Various applications of PAW such as anti-cancer ^7^, antimicrobial ^12^, anti-viral ^13^, food preservation ^5, 14^, insecticidal ^15^, oral hygiene ^16, 17^ and anti-fungal ^18^ have been demonstrated by various researchers. These studies have mostly worked with acidic pH PAW ^10, 18, 19^. However, the acidic pH of a solution can contribute to its corrosiveness, limiting its practical use, as prolonged exposure may accelerate the degradation of metals like stainless steel ^20^.

Real-world application of PAW requires the generation of neutral pH PAW, which has been successfully designed by our group before ^21, 22^. In this study, we highlight the bactericidal efficacy and mode of action of neutral-pH hs-PAbW (pH 7–7.4) against ESKAPE pathogens, which can cause pneumonia, surgical site, and bloodstream infections ^23, 24^. Demonstrating that hs-PAbW retains bactericidal activity at neutral pH enhances its physiological relevance, supporting its application in metal surface disinfection and wound cleansing to prevent hospital-acquired ESKAPE infections.

## Materials and Methods

### Generation of hs-PAbW, ROS-PAW, RNS-PAbW

We have generated hs-PAbW, ROS-PAW, RNS-PAbW by the protocol as described before. To maintain parity with our previous study, we have used the nomenclature hs-PAbW and RNS-PAbW as both have been generated using buffered water ^21^.

Briefly, hs-PAbW, ROS-PAW, and RNS-PAbW were generated using a 10 kV, 20 kHz AC plasma discharge in a 100 ml glass bottle with aluminium electrodes, treating 50 ml of liquid for 60 minutes, repeated at least three times. hs-PAbW, containing both ROS and RNS, was produced using 25 mM bicarbonate water with 1 LPM air bubbling and a 5 mm discharge, consuming 32 ± 2 W. ROS-PAW was generated from demineralized water with 1 LPM argon bubbling, a 5 mm discharge, and an ice jacket to prevent overheating, consuming 38 ± 2 W. RNS-PAbW was produced from 25 mM bicarbonate water with 1 LPM air bubbling and a 5 mm discharge, without an ice jacket, consuming 32 ± 2 W **(Figure 1A)**.

**Figure 1.**
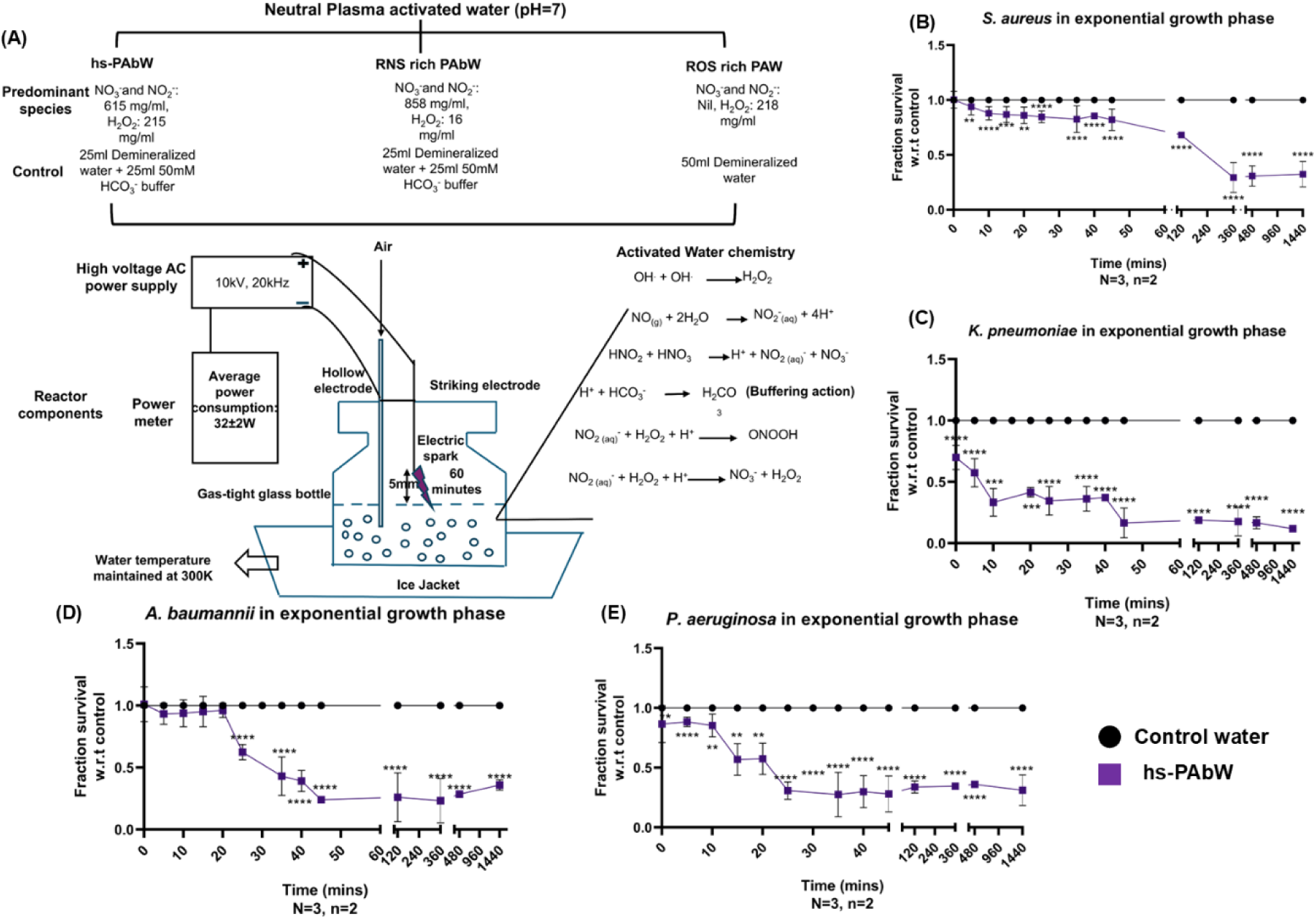
hs-PAbW induces a considerable amount of killing in exponentially growing bacteria. (A) Schematic depicting the generation of hs-PAbW, RNS-PAbW, and ROS-PAW. Line graphs illustrate fraction survival relative to the control treatment over exposure times ranging from 0 to 1440 minutes for *S. aureus* (B), *K. pneumoniae* (C), *A. baumannii* (D), and *P. aeruginosa* (E). The data are represented as mean ± SD of data points from 3 independent experiments. (N=3, n=2,). ***(P)* *< 0.05, *(P)* **< 0.005, *(P)* ***< 0.0005, *(P)* ****< 0.0001, ns= non-significant, (Unpaired two-tailed student’s t-test).**

### Bacterial strains and sample preparation

Methicillin Resistant *Staphylococcus aureus* (Isolate-SA3), hyper viscous (hv) *Klebsiella pneumoniae* (pus isolate 529) ^22^ (*S. aureus* and *K. pneumoniae* were kind gifts from Prof. Pradyut Prakash, Benaras Hindu University, India)^25^*, Acinetobacter baumannii* strain S2 (ATCC), and *Pseudomonas aeruginosa* PAO1 (a generous gift from Prof. Anne-Béatrice Blanc-Potard, Université de Montpellier) were revived on Luria-Bertani (LB) agar plates without antibiotics from glycerol stock stored at -80^0^ C. One colony was inoculated in the LB broth and grown overnight in a shaking incubator (170 rpm) at 37^0^C. After 12 hours, the stationary phase bacteria were subcultured (1:100) into freshly prepared LB broth and incubated for 4 to 5 hours to obtain the exponential growth phase. The (OD)_600_ of the bacterial cultures were adjusted to 0.3 (∼10^8^ CFU/ mL) and 0.1 (∼10^6^ CFU/ mL), respectively, centrifuged and the pellet was resuspended in 1 ml of control water, hs-PAbW, ROS-PAW, RNS-PAbW, and antibiotic solutions with high concentration (50 µg/ mL) of ceftazidime, ciprofloxacin, and meropenem for 45 minutes.

### Bacterial spread plating

Exponential-phase bacterial pellets of *S. aureus*, *K. pneumoniae*, *P. aeruginosa*, and *A. baumannii* were resuspended in 1 ml of water samples and incubated until the designated time points. After incubation, suspensions were serially diluted in 1X PBS, plated on LB agar, and incubated overnight at 37°C. Colony-forming units (CFU) were then recorded.

### Flow cytometry analysis

Bacterial pellets from exponential-phase cultures of *S. aureus*, *K. pneumoniae*, *A. baumannii*, and *P. aeruginosa* were incubated in 1 ml of control water, hs-PAbW, ROS-PAW, RNS-PAbW, or an antibiotic solution for 45 minutes. After centrifugation (6000 rpm, 10 min) and washing, cells were resuspended in 1 ml of double-autoclaved milli-Q water. To measure reactive nitrogen species, diaminofluorescein-2 diacetate (DAF_2_DA, 10 µM) was added and incubated for 30 minutes (excitation: 491 nm, emission: 513 nm) ^26^. Similarly, treated cells were incubated with dichlorodihydrofluorescein diacetate (DCF_2_DA, 10 µM) for reactive oxygen species, propidium iodide (PI, 1 µg/mL) for bacterial death, and DiBAC_4_ (1 µg/mL) for membrane depolarization. Samples were analysed via flow cytometry (BD FACSVerse, BD Biosciences), and data were processed using BD FACSuite software ^27^. Representative dot plots indicate the percentage of positive values.

### Scanning Electron Microscopy

Bacterial pellets from exponential cultures of *S. aureus*, *K. pneumoniae*, *P. aeruginosa*, *A. baumannii* were resuspended in 1 ml of the water samples. Post incubation of 45 mins, they were drop-casted on clear glass coverslips and air-dried. The samples were fixed by using 3.5% glutaraldehyde overnight at 4°C. The next day, they underwent a sequential dehydration process using 10%, 30%, 50%, 70%, 90%, and 100% alcohol, with each treatment lasting 3 minutes. The samples were dried using a vacuum-desiccator and subjected to gold spluttering before imaging. Thermofischer XL-30 ESEM was used for all the image acquisition at a magnification of 5000X. Bacteria on the metal surface were visualized using JEOL SEM IT-300 at the same magnification.

### Peroxynitrite Estimation Assay

The intracellular peroxynitrite assay was performed using Peroxynitrite Assay Kit from Abcam (ab2334690), using manufacturer’s protocol. Briefly, cell pellets from the exponential phase cultures of the tested pathogens were incubated in 1 ml of all the water samples for 45 minutes. Following this, the cells were centrifuged at 6,000 rpm for 10 minutes and the pellet was resuspended in fresh milliQ. 50 µl of the cell suspension was incubated with 50 µl of the reaction mix for 20 minutes. Following this, the fluorescence intensities were measured using a Tecan microplate reader, with Ex/Em at 490/530 nm.

### Atomic Force Microscopy

Bacterial pellets from exponential phase cultures of *S. aureus* and *K. pneumoniae* were resuspended in 1 ml of each water sample. After 45 minutes of incubation, the bacterial suspensions were centrifuged again at 6,000 rpm for 10 minutes, and the pellets were fixed using 3.5% paraformaldehyde. The bacterial suspensions were diluted 100-fold and 50 µl aliquot of each diluted sample was drop-cast onto coverslips and allowed to air dry. The images were captured using the NX10 AFM, and the length, height, and Young’s modulus of the bacterial cells were measured with XEI software^28^.

### Nitrotyrosine estimation

The pellets from exponential phase cultures of *S. aureus* and *K. pneumoniae* were centrifuged at 6,000 rpm for 10 minutes were resuspended in 1 ml of each water sample. After 45 minutes of incubation, the bacterial cells were lysed via sonication at 30% amplitude with 20-second pulses for a total of 4 minutes. Cell debris was removed by centrifugation at 13,000 rpm for 10 minutes at 4°C. The cell lysate was then concentrated using a speed vac, and protein concentration was determined using the Bradford reagent. An equal amount of protein was used for nitrotyrosine estimation, following the manufacturer’s instructions for the Nitrotyrosine ELISA Kit (ab210603) from Abcam. The end product of ELISA was quantified using a Tecan plate reader at 450 nm.

### Surface sterilisation experiment

Sterile stainless-steel equipment was obtained, and 10⁷ CFU of *S. aureus* and *K. pneumoniae* cultures were drop-casted onto the surfaces. After drying for one hour, 200 µl of RNS-PAbW and hs-PAbW were applied to the dried patches and incubated for an additional hour. The water samples were then removed, and the bacteria were resuspended in equal volumes of PBS before being spread plated to determine the CFU count post-exposure. For SEM-assisted bacterial visualization, the samples were fixed with 3.5% glutaraldehyde overnight at 4°C immediately after water removal. This was followed by dehydration, desiccation, and gold sputtering before imaging, as previously described.

### MTT Assay

The cytotoxicity of hs-PAbW was assessed using the MTT assay. In brief, L929 mouse fibroblast cells were seeded at a density of 5 × 10⁴ cells per well in a 96-well plate. Once the cells reached 70–80% confluency, they were exposed to varying concentrations of hs-PAbW in a total medium volume of 100 μl and incubated for 24 hours. Following the incubation, the cells were washed with 1× PBS, after which 100 μl of MTT solution (0.5 mg/mL) was added. The plate was then incubated for 4 hours at 37 °C in a CO₂ incubator. Afterward, the cells were treated with 50 μl of dimethyl sulfoxide (DMSO) to dissolve the formazan crystals formed by metabolically active cells. Absorbance was then recorded at 570 nm using a TECAN microplate reader.

### Ethics statement for animal experiments

All animal experiments were approved by the Institutional Animal Ethics Committee (IAEC) of the Indian Institute of Science, Bangalore (Registration No. 48/1999/CPCSEA). All procedures were conducted in strict accordance with the guidelines established by the Committee for the Purpose of Control and Supervision of Experiments on Animals (CPCSEA). The ethical clearance number for this study is CAF/Ethics/973/2023.

### Wound irrigation experiment

Wound irrigation experiments were performed by adapting previously established protocols ^29–31^. Briefly, 10–12-week-old male BALB/c mice were anesthetized intravenously on day 0 using ketamine hydrochloride (120 mg/kg). After dorsal depilation, two full-thickness excisional wounds (10 mm diameter) were created bilaterally along the midline using a sterile biopsy punch. Each wound was inoculated with 10^4^ CFU of *S. aureus* and allowed to incubate for 30 minutes before irrigation.

Each wound received 500 µL of the irrigation solutions, applied three times consecutively. Mice were housed individually post-surgery with access to standard diet. Wound irrigation continued alternate days until day 7, with wound images captured throughout the treatment period. On day 7, mice were euthanized, and wound tissues were harvested for bacterial quantification via spread plating on Mannitol Salt Agar (HiMedia). Wound contraction was assessed by measuring wound diameter using ImageJ software.

### Statistical Analysis

The number of biological and technical replicates used in each experiment have been reported in the figure legends. The statistical analyses were done by unpaired two-tailed student’s t-test, one-way ANOVA, or Mann-Whitney U-test as indicated in the figure legends. The *p* values below 0.05 were considered significant. The results were expressed as mean ± SD or mean± SEM as mentioned in the figure legends. GraphPad Prism 8.4.3 (686) was used to generate the bar graphs and perform the statistical tests.

## Results

### Plasma-activated water with a neutral pH could eliminate bacteria in the exponential growth phase

We subjected exponential-phase cultures of *Staphylococcus aureus*, *Klebsiella pneumoniae*, *Acinetobacter baumannii*, and *Pseudomonas aeruginosa* to hs-PAbW treatment to evaluate its bactericidal efficacy against these ESKAPE pathogens and determine the minimum exposure time required for its action **(Figure 1B–E)**. hs-PAbW reduced bacterial survival across all species, with varying minimum exposure times: *S. aureus* showed a significant reduction from 5 minutes onwards **(Figure 1B)**, *A. baumannii* from 25 minutes onwards **(Figure 1D)**, and hypervirulent *K. pneumoniae* and *P. aeruginosa* started showing immediately **(Figure 1C, E)**. A 45-minute treatment, which significantly reduced viability across all species, was chosen for further experiments. *A. baumannii* and *P. aeruginosa* reached maximum reduction at this point **(Figure 1D, E)**. Survival of *K. pneumoniae* **(Figure 1C)** continued to decrease until 1440 minutes, and while *S. aureus* **(Figure 1B)** also showed further reduction, substantial killing was achieved by 45 minutes.

To determine whether hs-PAbW also exhibited a bactericidal effect on stationary-phase bacteria, we treated stationary-phase cultures of *S. aureus*, *K. pneumoniae*, *A. baumannii*, and *P. aeruginosa* with hs-PAbW for 45 minutes. The absence of any change in the percentage of propidium iodide (PI)-positive cells indicated that hs-PAbW had no effect on the viability of stationary-phase bacterial cultures **(Figure S1A-S1H)**. From this point forward, exponential-phase cultures were used for all experiments.

### Unlike the nitrogen species (NO_2_^-^, NO_3_^-^) present in hs-PAbW, oxygen species (H₂O₂) alone is not capable of independently killing bacterial pathogens

The microbicidal activity of hs-PAbW can be attributed to its high content of ROS and RNS. We sought to identify the role of each of these functional components in reducing bacterial growth. As we found that hs-PAbW can restrict bacterial growth at the exponential phase, we subjected the exponential phase cultures of all three bacterial strains to the treatment with ROS-PAW **(Figure 2)** and RNS-PAbW **(Figure 3)**.

**Figure 2.**
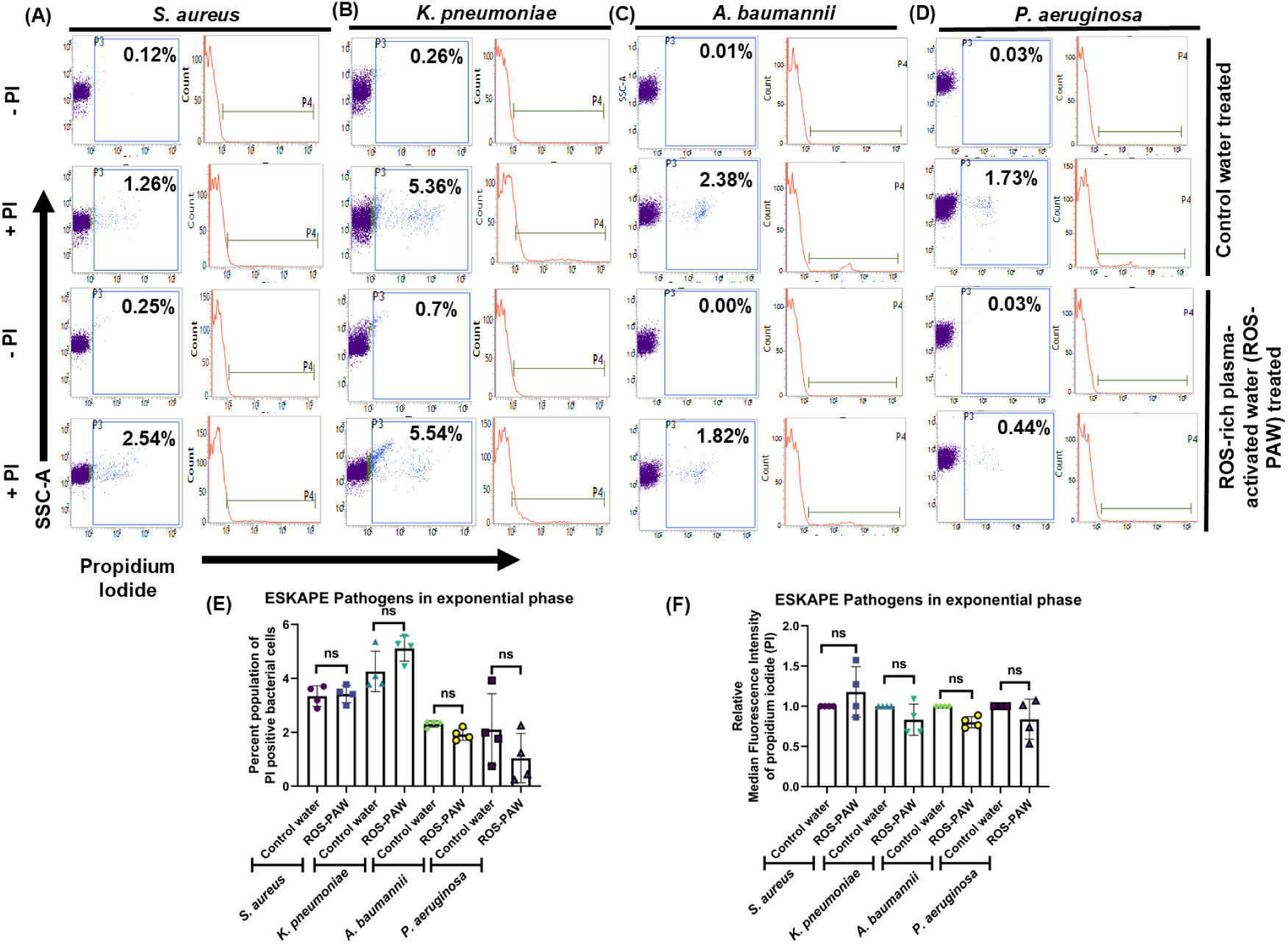
ROS-PAW itself is unable to induce bacterial death. (A, B, C, and D) The dot plot (SSC-A vs. PI) and histogram (count vs. PI) represent the viability of exponentially growing *S. aureus* (A), *K. pneumoniae* (B), *A. baumannii* (C), and *P. aeruginosa* (D) treated with reactive oxygen species rich plasma-activated water (ROS-PAW). (D) The bar graphs represent the PI-positive percent population of *S. aureus*, *K. pneumoniae*, *A. baumannii*, and *P. aeruginosa* with or without ROS-PAW treatment. (E) The median fluorescence intensity (MFI) of the PI-positive population of *S. aureus*, *K. pneumoniae*, *A. baumannii*, and *P. aeruginosa* with or without ROS-PAW treatment (F). The data are representative of n≥3, N=3 and are presented as mean ± SD ***(P)* *< 0.05, *(P)* **< 0.005, *(P)* ***< 0.0005, *(P)* ****< 0.0001, ns= non-significant, (Unpaired two-tailed student’s t-test).**

**Figure 3.**
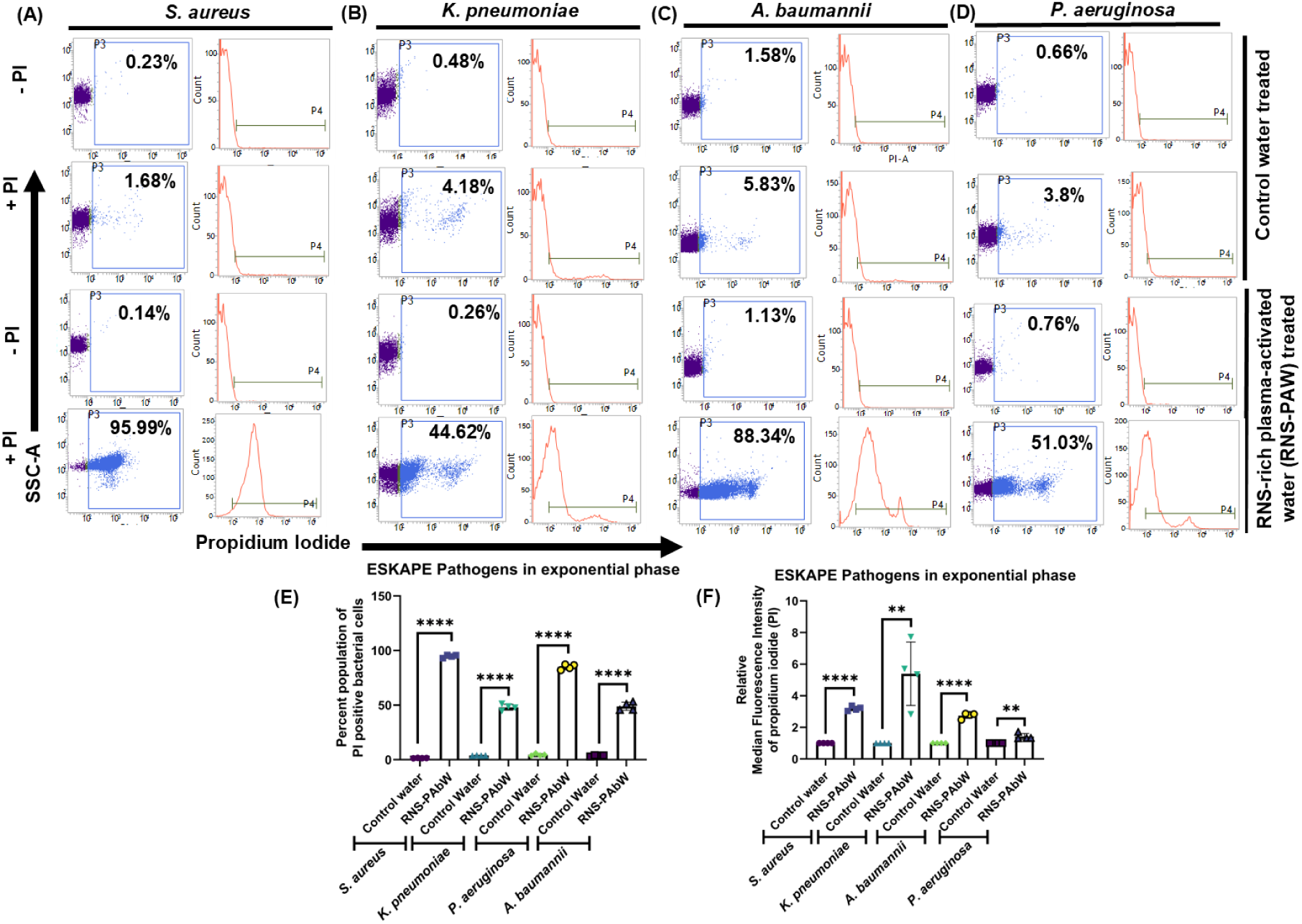
RNS-PAbW can independently kill exponentially growing bacteria. (A, B, C, and D) The dot plot (SSC-A vs. PI) and histogram (count vs. PI) represent the viability of exponentially growing *S. aureus* (A), *K. pneumoniae* (B), *A. baumannii* (C), and *P. aeruginosa* (D) treated with reactive nitrogen species rich plasma-activated water (RNS-PAbW). The bar graphs represent the PI-positive percent population of *S. aureus*, *K. pneumoniae*, *A. baumannii*, and *P. aeruginosa* with or without RNS-PAbW treatment. (E) The relative median fluorescence intensity (MFI) of the PI-positive population of *S. aureus*, *K. pneumoniae*, *A. baumannii*, and *P. aeruginosa* with or without RNS-PAbW treatment (F). The data are representative of n≥3, N=3 and are presented as mean ± SD ***(P)* *< 0.05, *(P)* **< 0.005, *(P)* ***< 0.0005, *(P)* ****< 0.0001, ns= non-significant, (Unpaired two-tailed student’s t-test).**

ROS-PAW and RNS-PAbW were prepared to contain ROS and RNS concentrations equivalent to those in hs-PAbW **(Figure 1A)**. ROS-PAW treatment resulted in a higher DCF_2_DA-median fluorescence intensity (MFI), indicating elevated ROS levels **(Figure S2A-E).** However, PI staining showed that ROS-PAW was ineffective at killing bacteria despite high intracellular ROS levels **(Figure 2A-F)**. The greater DAF_2_DA-MFI value **(Figure S3A-E)** corresponding to RNS-PAbW-treated bacterial cells further showed that treating the bacterial cells with RNS-PAbW generated high concentration of RNS. Unlike ROS-PAW, we observed RNS-PAbW could kill the actively growing bacteria **(Figure 3A-3F)**. Bacterial spread plating confirmed these findings **(Figure S4 A-D)**. DAF_2_DA staining of hs-PAbW-treated exponential **(Figure S5A-H)** cultures of the test pathogens also showed significantly higher amounts of RNS compared to the control water treatment. Taken together, we could conclude that RNS is the major factor contributing to the microbicidal activity of hs-PAbW (pH=7 to 7.4) in ESKAPE pathogens.

### High concentration of peroxynitrite in RNS-PAbW and hs-PAbW is one of the factors regulating the microbicidal role of hs-PAbW

Morphological changes were observed in *K. pneumoniae*, *A. baumannii*, and *P. aeruginosa* after hs-PAbW treatment compared to the control **(Figure 4A)**. *K. pneumoniae* exhibited shrinkage, distortion, and clumping, while *A. baumannii* showed visible lysis, and *P. aeruginosa* displayed reduced cell size and clumping. ROS-PAW and RNS-PAbW also induced surface morphology changes in all three strains, as seen via SEM. In contrast*, S. aureus* showed no noticeable morphological alterations **(Figure S6)**.

**Figure 4.**
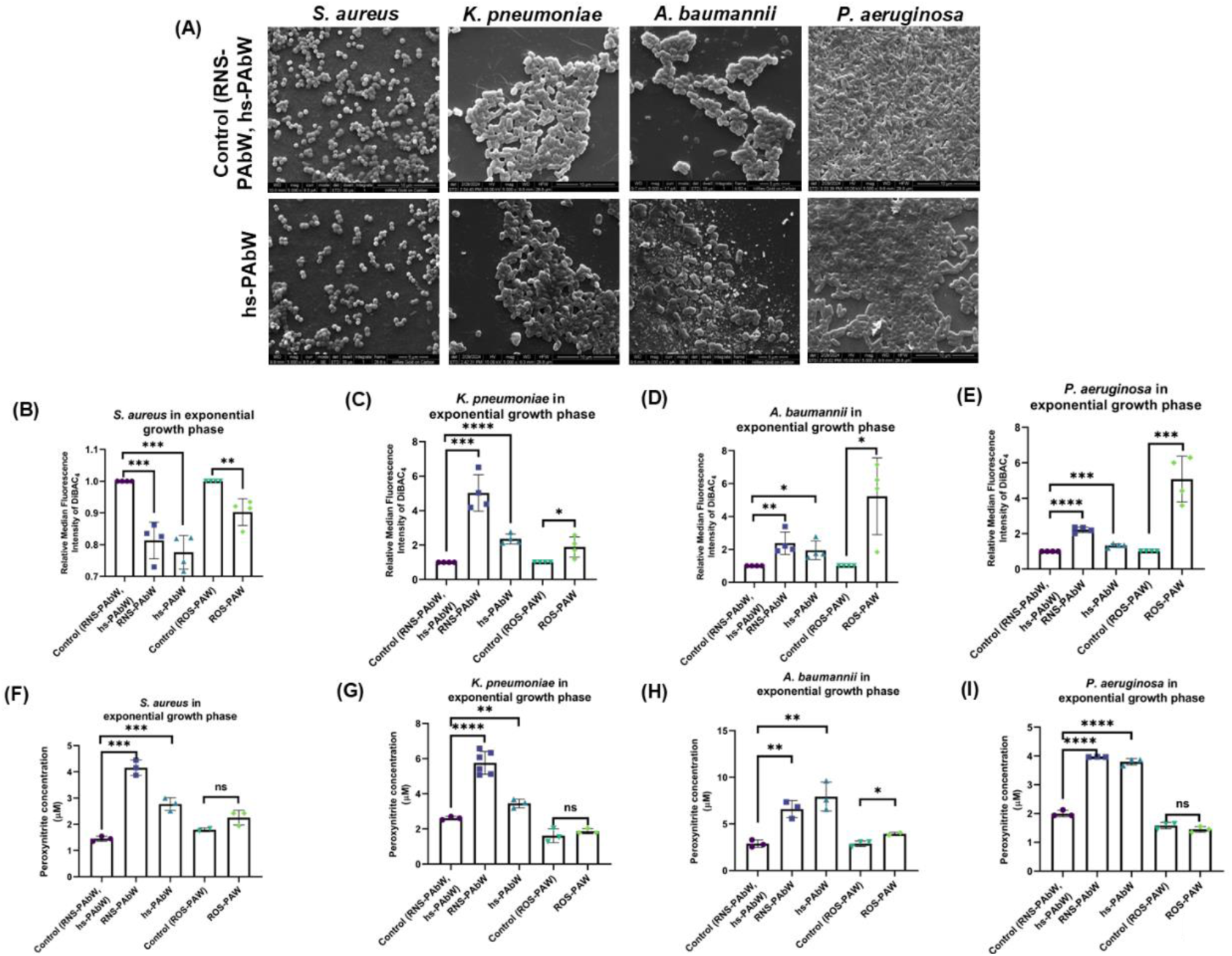
ROS-PAW can alter membrane potential, however, it is not sufficient for a bactericidal effect. Scanning Electron Microscopy images of *S. aureus*, *K. pneumoniae*, *A. baumannii*, *P. aeruginosa* treated with hs-PAbW and control water (5000X magnification) (A). The relative median fluorescence intensity (MFI) of the DiBAC_4_-positive population of *S. aureus*, *K. pneumoniae*, *A. baumannii*, and *P. aeruginosa* with RNS-PAbW, hs-PAbW, ROS-PAW treatment. is represented in figure panels B-E. The data are representative of n≥3, N=3 and are presented as mean ± SD. Figure panels **F-I** contain the bar graphs that represent the quantity of intracellular peroxynitrite produced upon treatment with control ROS-PAW, ROS-PAW, control RNS-PAbW, RNS-PAbW, control hs-PAbW and hs-PAbW of *S. aureus* (B), *K. pneumoniae* (C), *A. baumannii* (D), *P. aeruginosa* (E). The data are representative of n≥3, N=3 and are presented as mean ± SD ***(P)* *< 0.05, *(P)* **< 0.005, *(P)* ***< 0.0005, *(P)* ****< 0.0001, ns= non-significant, (Unpaired two-tailed student’s t-test).**

Previous studies on *Fusarium graminearum* spores suggested PAW compromises membrane integrity, leading to intracellular content leakage^32^. To investigate this in our samples, we used the membrane-potential-sensitive dye DiBAC_4_. Results indicated increased membrane depolarization in *P. aeruginosa*, *K. pneumoniae*, and *A. baumannii* **(Figure 4C-E)**, whereas *S. aureus* exhibited membrane hyperpolarization **(Figure 4B)**. Although ROS-PAW and RNS-PAbW altered membrane potential, only hs-PAbW demonstrated bactericidal effects, suggesting membrane potential disruption alone is not responsible for its antimicrobial action.

We aimed to identify the key chemical species responsible for the bactericidal activity of neutral pH PAW. A significant increase in the PI positive population after RNS-PAbW treatment suggests that RNS in hs-PAbW contribute to its microbicidal effect against ESKAPE pathogens. In addition to ROS and RNS, hs-PAbW contains short-lived species like peroxynitrous acid (ONOOH) and peroxynitrite (ONOO^-^), which can cause cell damage through direct oxidation or decomposition byproducts^6, 8, 21, 33^. To assess their role, we measured intracellular peroxynitrite levels post-treatment **(Figure 4F-4I)**. While ROS-PAW showed no significant change, RNS-PAbW and hs-PAbW treatments led to a notable increase. In *S. aureus* and *K. pneumoniae* exponential phase culture, RNS-PAbW produced significantly higher peroxynitrite levels than hs-PAbW **(Figure 4F-G)**, whereas in *A. baumannii* and *P. aeruginosa*, both treatments generated comparable peroxynitrite levels **(Figure 4H-I)**. This suggests that RNS in hs-PAbW regulate peroxynitrite formation.

### RNS-PAbW, hs-PAbW, and ROS-PAbW caused a reduction in the length, height, and Young’s modulus of the Gram-negative bacterium *K. pneumoniae* although ROS-PAW was unrelated to increased nitrotyrosine levels in *S. aureus* and *K. pneumoniae*

We selected *S. aureus* and *K. pneumoniae* as representative Gram-positive and Gram-negative bacteria, respectively, and analysed their morphological changes using Atomic Force Microscopy **(Figure 5A-F)**. Consistent with SEM data, water sample treatments did not significantly affect *S. aureus* in terms of diameter or Young’s modulus **(Figure 5B, C)**. However, *K. pneumoniae* showed a significant reduction in these parameters after exposure to RNS-PAbW, hs-PAbW, and ROS-PAW **(Figure 5D-F).** This suggests that while ROS-PAW does not effectively kill bacteria, it can impact structural integrity, reinforcing that RNS and short-lived species like peroxynitrite, rather than ROS, drive the bactericidal effects of hs-PAbW.

**Figure 5.**
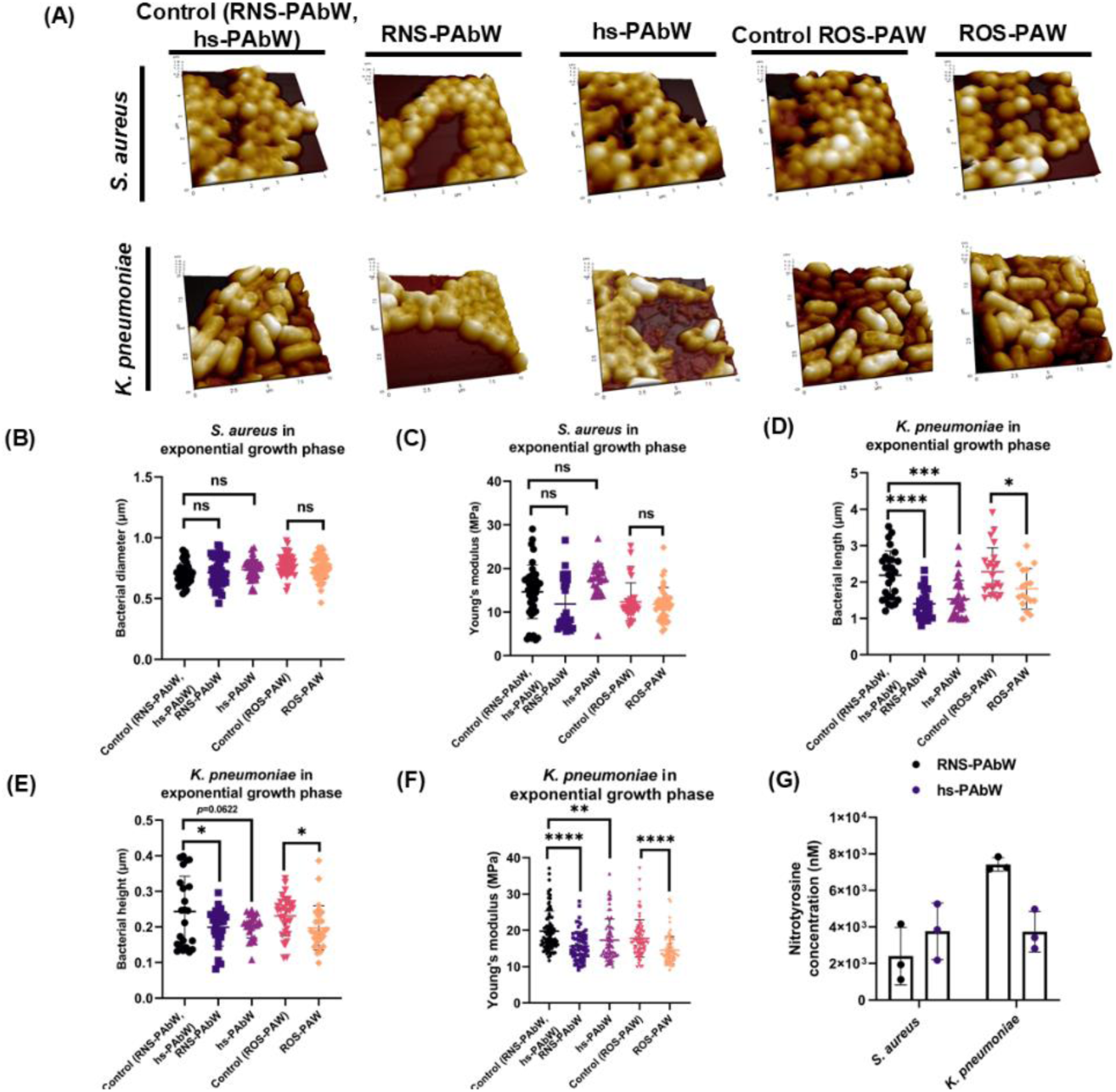
Treatment with RNS-PAbW, hs-PAbW, and ROS-PAbW caused a reduction in the length, height, and Young’s modulus of the Gram-negative bacterium *K. pneumoniae* although ROS-PAW was unrelated to increased nitrotyrosine levels in *S. aureus* and *K. pneumoniae*. (A) Atomic Force Microscopy images of *S. aureus* and *K. pneumoniae* after treatment with control waters, ROS-PAW, hs-PAbW, and RNS-PAbW. Quantification of the length, height, and Young’s modulus for *S. aureus* (B, C, D) and *K. pneumoniae* (E, F, G) was performed. Data for (A-G) are representative data of N=2, n≥20, and are shown as mean±SD. (H) shows intracellular nitrotyrosine levels in *S. aureus* and *K. pneumoniae* following treatment with RNS-PAbW and hs-PAbW. The data is a representative of N=2, n=3, and is presented as mean±SD. *(P)* *< 0.05, *(P)* **< 0.005, *(P)* ***< 0.0005, *(P)* ****< 0.0001, ns= non-significant, (Unpaired two-tailed student’s t-test).

Peroxynitrous acid can nitrate both free and protein-bound tyrosine, with 3-nitrotyrosine serving as a biomarker for peroxynitrite formation ^34, 35^. This modification can disrupt protein structure and function and is a key defence mechanism in macrophages ^36, 37^. In our study, nitrotyrosine was significantly detected in *S. aureus* and *K. pneumoniae* cell lysates after hs-PAbW and RNS-PAbW exposure, indicating active peroxynitrite formation and its antimicrobial role **(Figure 5H)**. As expected, nitrotyrosine was undetectable in bacteria treated with control water and ROS-PAW, reinforcing that RNS and peroxynitrite are crucial for the bactericidal activity of neutral pH hs-PAbW. The standard curve and raw data are provided in supplementary data **(Figure S7)**.

### hs-PAbW and RNS-PAbW efficiently reduce microbial load on metal surfaces, supporting their use as surface disinfectants

Hospital surfaces such as floors and door handles serve as key reservoirs for nosocomial infections, increasing the risk of infection spread both within and beyond the hospital^38, 39^. Given the antimicrobial potential of hs-PAbW observed in our previous data, we aimed to explore its potential use as a surface cleansing agent. We used sterile metal surfaces and drop-casted *S. aureus* and *K. pneumoniae* cultures onto them. Both RNS-PAbW and hs-PAbW effectively reduced the bacterial load on metal surfaces after one hour of exposure, as verified by bacterial spread plating **(Figure 6A, B)** and SEM-assisted imaging of infected metal surfaces **(Figure 6C)**, underscoring their potential as a cost-effective surface decontamination solution.

**Figure 6.**
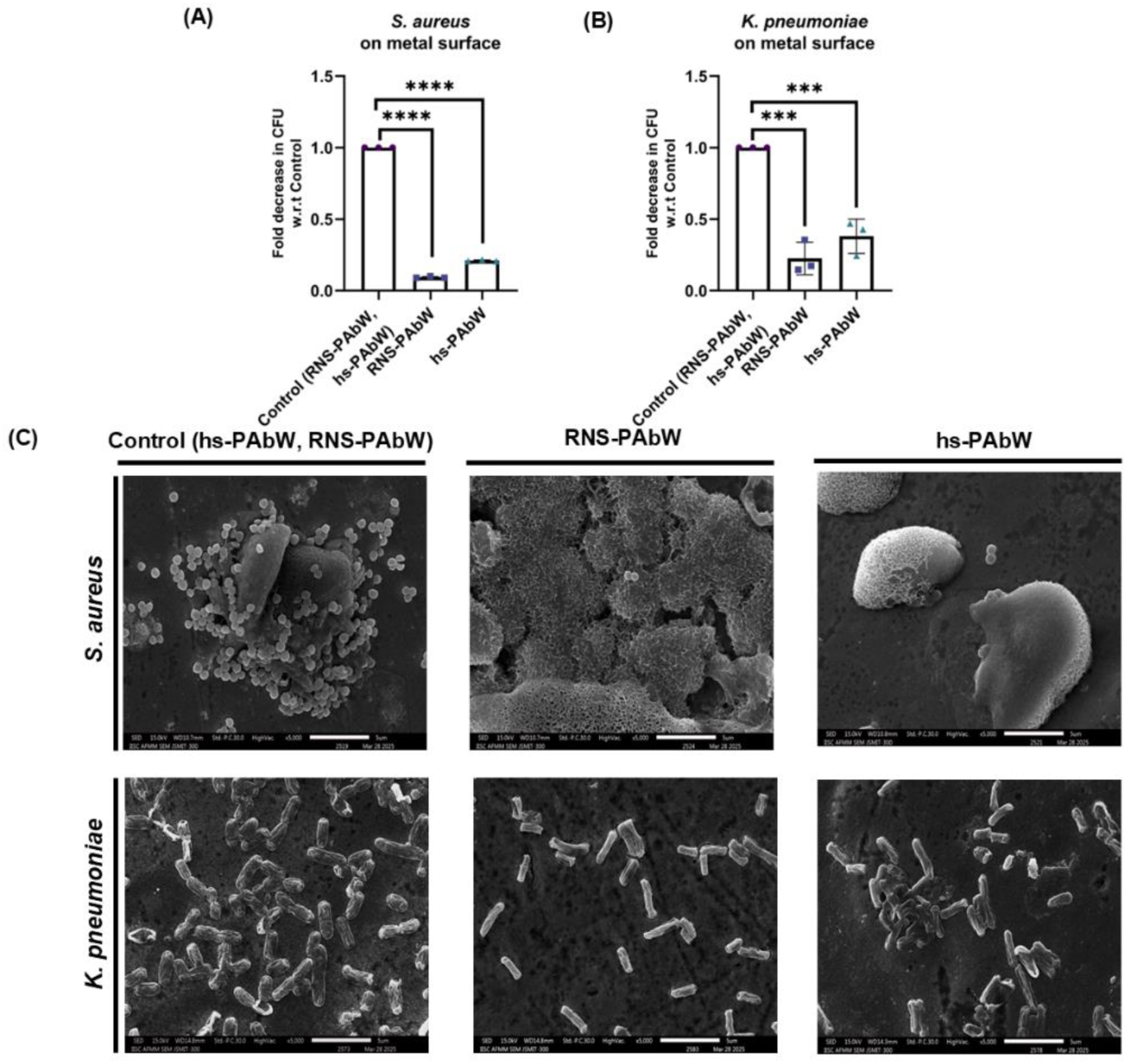
hs-PAbW can successfully reduce bacterial burden on non-autoclavable metal surfaces. Bar graphs (A) and (B) depict the fold reduction in CFU after treatment with hs-PAbW and RNS-PAbW on *S. aureus* (A) and *K. pneumoniae* (B) droplets dried on metal surfaces. The data are represented as mean ± SEM. (N=3, n=2). Figure panel (G) includes SEM images of *S. aureus* and *K. pneumoniae* on stainless steel surface after cleansing with control water, hs-PAbW, RNS-PAbW (5000X magnification). ***(P)* *< 0.05, *(P)* **< 0.005, *(P)* ***< 0.0005, *(P)* ****< 0.0001, ns= non-significant, (Unpaired two-tailed student’s t-test).**

### Both hs-PAbW demonstrated superior efficacy against ESKAPE pathogens *in vitro* compared to high-dose antibiotics and effectively reduced bacterial burden in infected murine wounds, outperforming commercially available antibiotic

Ceftazidime (third-generation cephalosporin) and meropenem (carbapenem) are β-lactam antibiotics that are routinely used in treating a wide range of MDR bacterial infections ^28, 40, 41^. Ciprofloxacin is a fluoroquinolone that inhibits the growth of several pathogens by blocking the activity of DNA gyrase (*gyrAB*) ^42–44^.Since hs-PAbW could successfully reduce the viability of actively growing bacteria, we decided to compare its bactericidal activity with these antibiotics. We harvested bacteria from the exponential phase of their growth and surprisingly, we observed that hs-PAbW is more effective in inducing bacterial death compared to high concentrations of antibiotics (ceftazidime, ciprofloxacin, and meropenem, concentration-50 µg/ mL) and hs-PAbW within 45 minutes **(Figure 7A-D)**. The reduced susceptibility of *K. pneumoniae* to meropenem can be overcome by hs-PAbW^45^.

**Figure 7:**
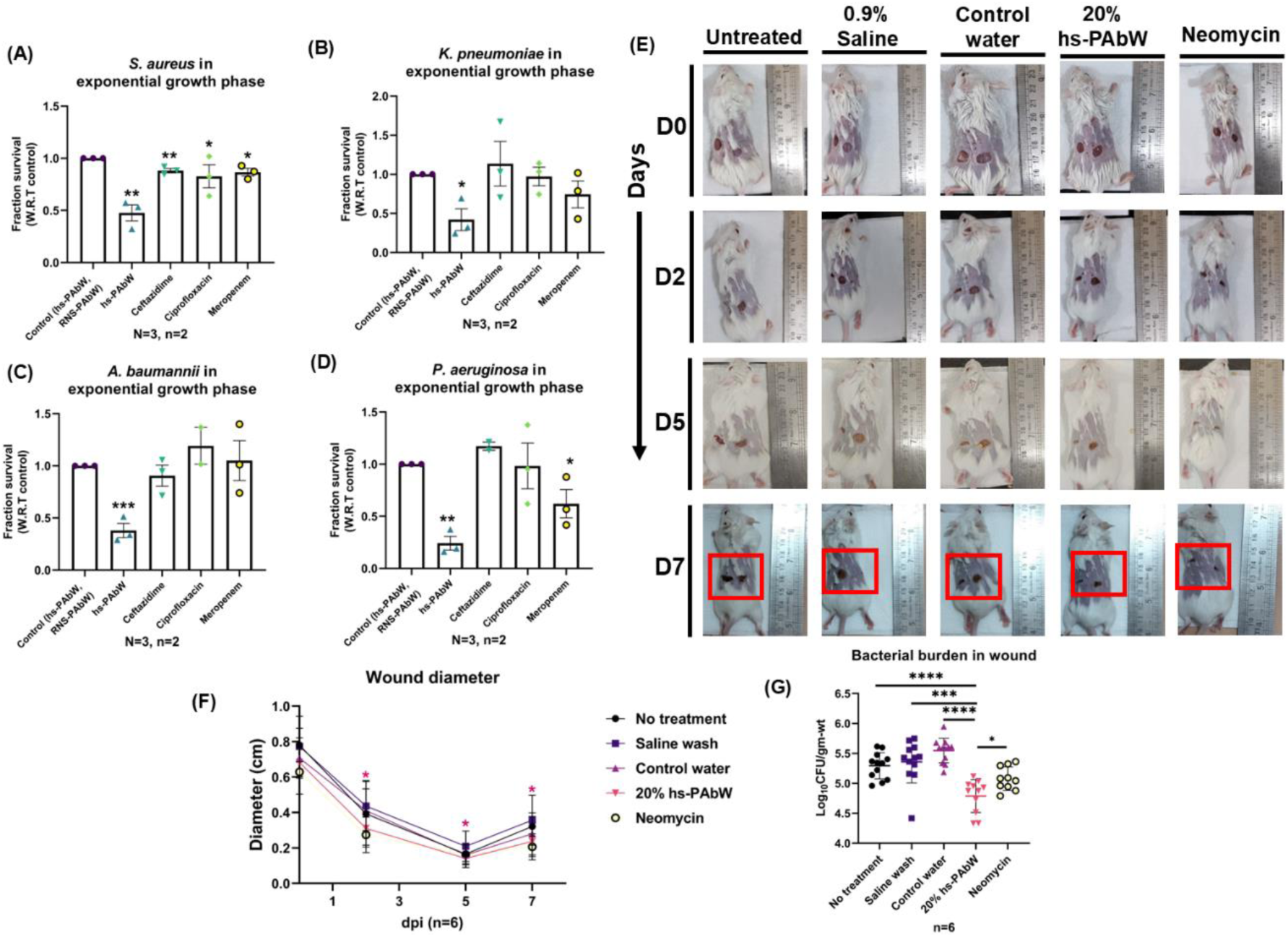
hs-PAbW was more effective at killing ESKAPE pathogens than high doses of antibiotics, and a non-cytotoxic dose of the same can be used as a wound irrigation agent. Bar graphs (A-D) depict the reduction in CFU of *S. aureus*, *K. pneumoniae*, *A. baumannii*, *P. aeruginosa* after treatment with hs-PAbW and 50 µg/ml of ceftazidime, meropenem, and ciprofloxacin *in-vitro*. The data are represented as mean ± SEM. (N=3, n=2). The images show wound closure in Balb/c mice infected with *S. aureus*, following treatment with various irrigation methods including 0.9% saline, control water, hs-PAbW, and neomycin. Wound diameter (F) and bacterial load (G) were quantified, with data presented as mean ± SD for n = 6. *(P)* *< 0.05, *(P)* **< 0.005, *(P)* ***< 0.0005, *(P)* ****< 0.0001, ns= non-significant, (Unpaired two-tailed student’s t-test) (Mann-Whitney U-test for *in-vivo* experiments).

To evaluate the suitability of hs-PAbW as a disinfectant, its cytotoxicity was assessed. No significant cytotoxicity was observed up to 20% concentration (**Fig. S8A**). Notably, hs-PAbW retained its antimicrobial activity even when diluted to 20%, effectively reducing bacterial counts in vitro (**Fig. S8B**). Irrigating the wound with this non-cytotoxic concentration of hs-PAbW led to faster wound closure compared to control treatments such as water rinse and 0.9% saline, showing comparable effects to topical Neomycin application (**Figure 7E, F**). This improvement was also accompanied by a marked reduction in wound bacterial burden (**Figure 7G**).

## Discussion

PAW, which contains various polyatomic ions such as NO_2_^−^, _3_^−^, and H_2_O_2_ (in this study: NO_2_^−^ at 600 mg/L, _3_^−^ at 130 mg/L, and H_2_O_2_ at 200 mg/L), has previously demonstrated significant potential in bacterial decontamination ^46, 47^. The combination of nitric oxide and superoxide ions in acidic pH gives rise to several secondary reactive nitrogen intermediates, such as peroxynitrite (ONOO^−^, ONOOH), which can function as nucleophilic oxidants ^48, 49^. Due to its high ROS and RNS content (RONS), PAW is widely used in food and agricultural industries for disinfecting fruits and raw vegetables ^11, 50^. Multiple studies showed that acidic pH stabilizes and intensifies the activity of reactive oxygen and nitrogen species ^51, 52^. However, generating PAW at neutral pH is required for real world applications. In one of our previous studies, we demonstrated that using bicarbonate water can effectively buffer and maintain the pH of PAW near neutrality (hs-PAbW), making it more practical for real-world applications. Additionally, we showed the antimicrobial effect of hs-PAbW on *K. pneumoniae* and methicillin-resistant *S. aureus* (MRSA) in the same study ^21^. However, the mechanism behind the bactericidal action of hs-PAbW remained unclear.

In this study we have seen that DCF_2_DA and DAF_2_DA staining of hs-PAbW, ROS-PAW, and RNS-PAbW treated bacterial samples showed that the stability of RONS was equally maintained in the physiological pH range (between 7 and 7.4). However, it was observed that hs-PAbW is more efficient in killing exponential-phase bacteria than the stationary phase. The high rate of metabolic activity and basal respiration could be a reason for the greater sensitivity of the exponential phase cultures to hs-PAbW ^53^. We also show that ROS-PAW, which contained similar levels of ROS as hs-PAbW, couldn’t kill the bacteria independently, proving that their concentrations in hs-PAbW are unable to bring about the bactericidal effect independently. Instead, when the bacterial strains are incubated in RNS-PAbW where RNS are present at a specific proportion (as mentioned earlier), they execute the maximum bactericidal activity. Incubation in ROS-PAW and RNS-PAbW resulted in changed membrane potential of all the bacterial strains tested, like hs-PAbW. As reported previously, this effect may be caused by lipid peroxidation mediated by reactive species present in PAW, which leads to oxidative degradation of cholesterol, phospholipids, and unsaturated fatty acids, ultimately causing damage to the cell membrane ^54–59^. Also, previous studies had shown that the thick cell wall in Gram-positive bacteria confers resistance to PAW. This can explain why we observed no change in the size or stiffness of *S. aureus* upon exposure to any of the water samples ^60^.

hs-PAbW exposure generates intracellular peroxynitrite, a key reactive nitrogen species (RNS) also abundant in RNS-PAbW-treated cells. Peroxynitrite induces oxidative and nitrosative stress by modifying proteins, oxidizing lipids, and damaging DNA, either directly or through its decomposition ^61–63^. Oxidative stress induced by peroxynitrite can result from the direct oxidation reactions of peroxynitrous acid or from the generation of oxidizing radicals (e.g., •OH). It leads to oxidation of thiols, heme proteins, and iron-sulfur clusters, as well as nitration of tyrosine residues, which can interfere with bacterial protein function ^36, 61, 64^. Due to negligible peroxynitrite formation upon ROS-PAW treatment, nitrotyrosine levels were also low in that condition. This shows that nitrotyrosine formation is one of the primary mechanisms responsible for the bactericidal action of our hs-PAbW. We also show that RNS in hs-PAbW is the key factor driving its bactericidal effect. Thus, we successfully eliminate the need for acidic pH for PAW activity and have developed neutral pH hs-PAbW, enhancing its relevance for real-world applications.

hs-PAbW effectively decontaminated non-autoclavable metal surfaces, presenting a cost-effective disinfection strategy for hospital environments. A 45-minute exposure to hs-PAbW resulted in significantly greater bacterial killing *in vitro* compared to high doses of antibiotics. Moreover, in a murine model of *S. aureus*-infected wounds, a non-cytotoxic dose of hs-PAbW promoted superior wound healing compared to 0.9% saline and comparable to topical neomycin treatment. We propose a single hs-PAbW setup in hospitals, where technicians can generate it via controlled electric discharges. This sustainable approach will require only buffered water, minimizing raw material transport, waste, and carbon footprint. Ethically sourced materials and the absence of harmful emissions will ensure safety. While hs-PAbW loses efficacy after seven days, reactive species degradation can be managed with inhibitors like vitamin C and isoflavones ^21, 65–68^. Further research is needed to assess its environmental safety post-inhibition.

To the best of our knowledge, this is the first study to highlight the mechanism of action of neutral pH hs-PAbW and discuss its potential to prevent spread of ESKAPE pathogens.

## Author contribution

Conceptualization: DM, ARC, LNR, and DC, Methodology: DM, PG, ARC, NV Investigation: DM, PG, ARC, NV Visualization: DM,ARC, Data curation: DM, PG, ARC, Formal analysis: DM, ARC, Fund acquisition: LNR and DC, Project administration: DM, ARC, LNR, and DC, Writing-original draft: DM, ARC, and LNR, Editing and proofreading: PG, and DC, Supervision: LNR and DC.

## Acknowledgments

The authors thank the central flow cytometry facility at IISc. Mrs. Dharani and Mr. Munish from the central flow cytometry facility are acknowledged for their help in FACS data acquisition. Mrs. Deepika from Advanced Facility for Microscopy and Microanalysis (AFMM), IISc is acknowledged for her help in acquiring Scanning Electron Microscopy (SEM) images. Mrs. Monisha from Atomic Force Microscopy (AFM) Facility, Department of Bioengineering (BE), IISc, is acknowledged for her help in acquiring the AFM images.

## Transparency statement

None to declare.

## Data availability

All data generated for this study have been included in the manuscript.

## Supplementary Figures

**Figure S1.**
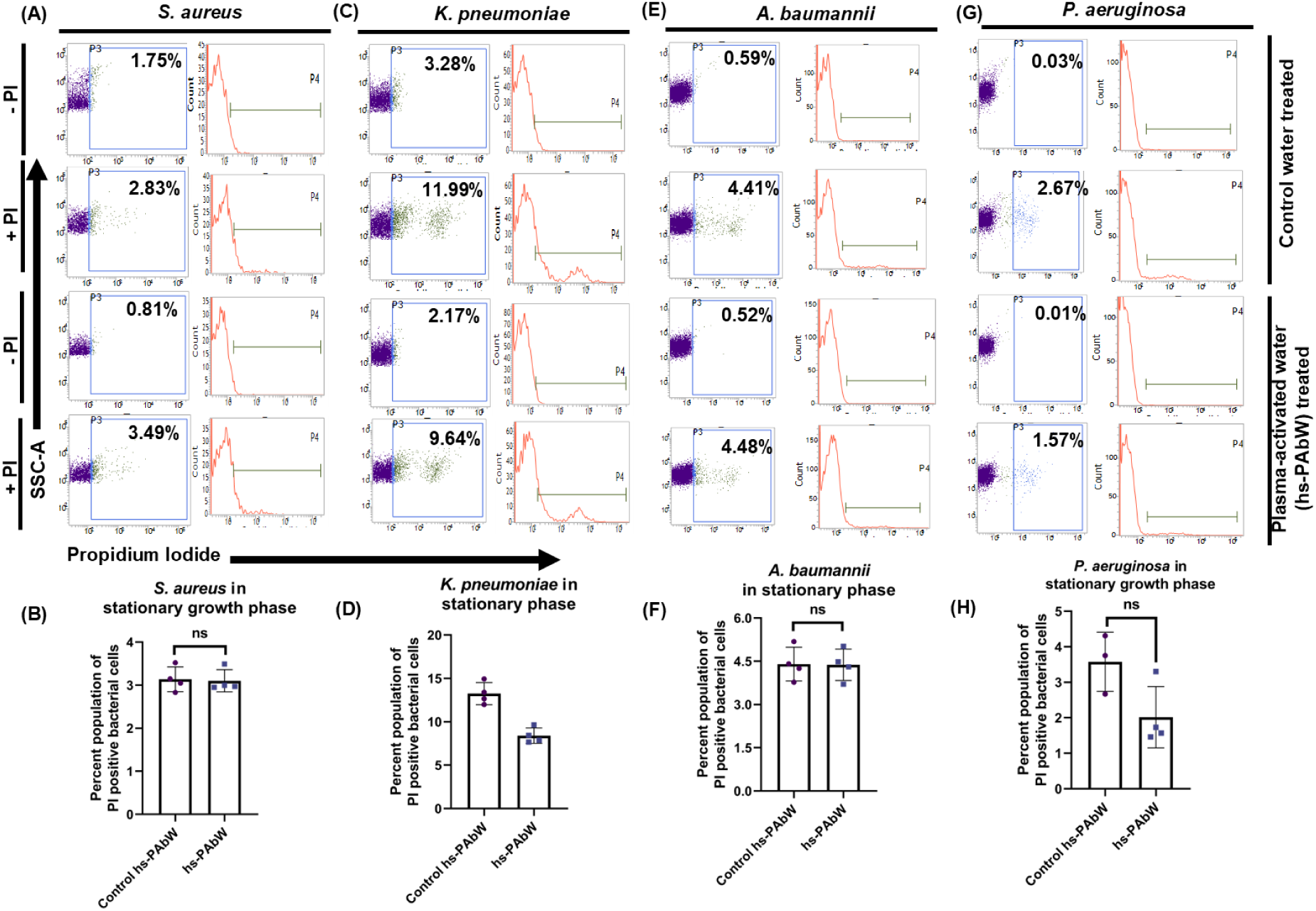
hs-PAbW couldn’t kill the stationary phase cultures of bacteria. (A, C, E and G) The dot plot (SSC-A vs. PI) and histogram (count vs. PI) represent the viability of exponentially growing cultures of *S. aureus* (A), *K. pneumoniae* (C), *A. baumannii* (E), and *P. aeruginosa* (E) with hs-PAbW. The bar graphs represent the propidium iodide positive percent population of *S. aureus* (B), *K. pneumoniae* (D), *A. baumannii* (F), and *P. aeruginosa* (H) with or without hs-PAbW treatment. The data are representative of n≥3, N=3 and is presented as mean ± SD. ***(P)* *< 0.05, *(P)* **< 0.005, *(P)* ***< 0.0005, *(P)* ****< 0.0001, ns= non-significant, (Unpaired two-tailed student’s t-test).**

**Figure S2.**
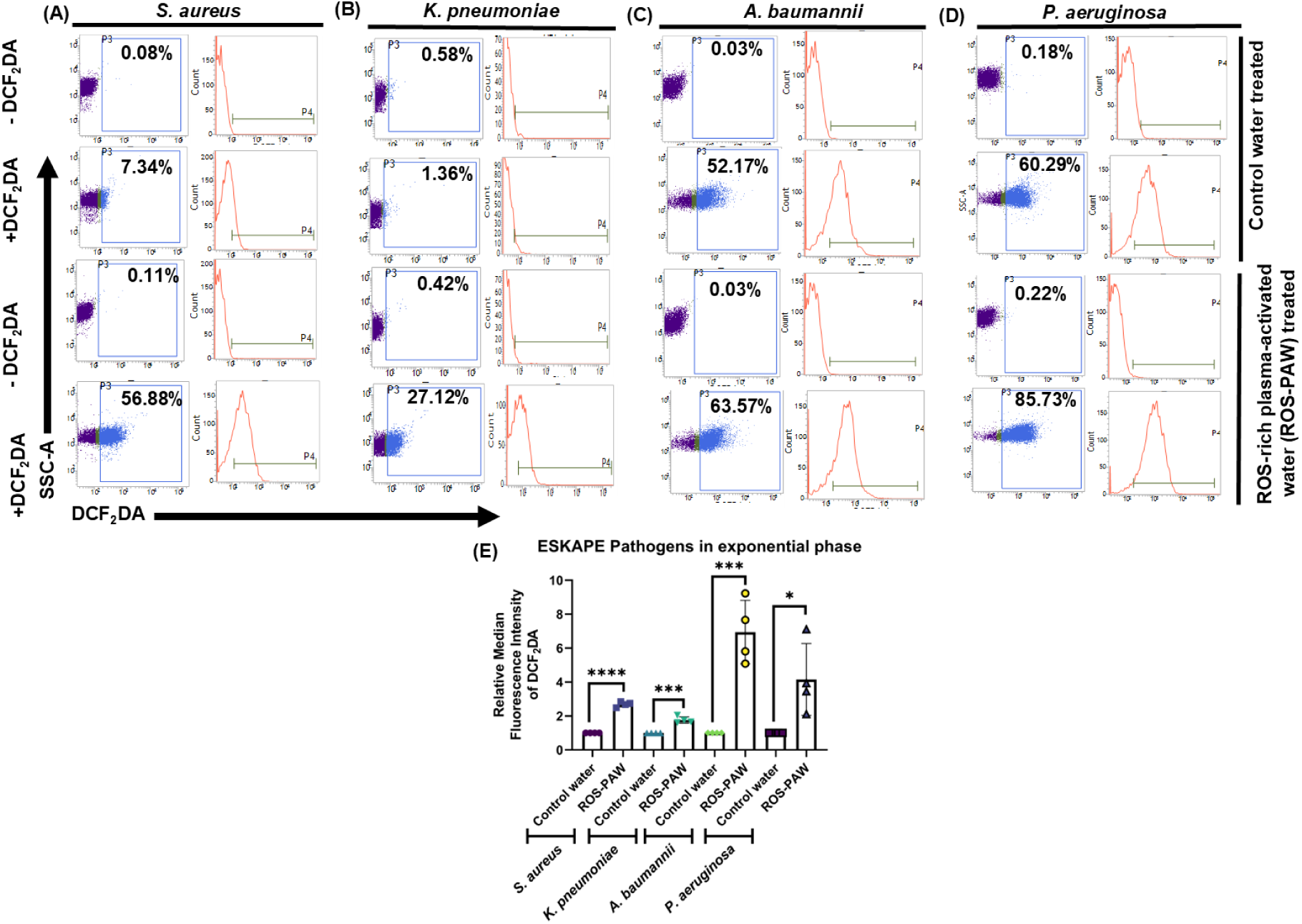
ROS-PAW generates a significant amount of ROS. (A, B, C, and D) The dot plot (SSC-A vs. DCF_2_DA) and histogram (count vs. DCF_2_DA) represent the ROS content of exponentially growing *S. aureus* (A), *K. pneumoniae* (B), *A. baumannii* (C), and *P. aeruginosa* (D) treated with reactive oxygen species rich plasma-activated water (ROS-PAW). The bar graphs represent the relative median fluorescence intensity (MFI) of the DCF_2_DA-positive population of *S. aureus*, *K. pneumoniae*, *A. baumannii*, and *P. aeruginosa* with or without ROS-PAW treatment (E). The data are representative of n≥3, N=3 and is presented as mean ± SD. ***(P)* *< 0.05, *(P)* **< 0.005, *(P)* ***< 0.0005, *(P)* ****< 0.0001, ns= non-significant, (Unpaired two-tailed student’s t-test).**

**Figure S3.**
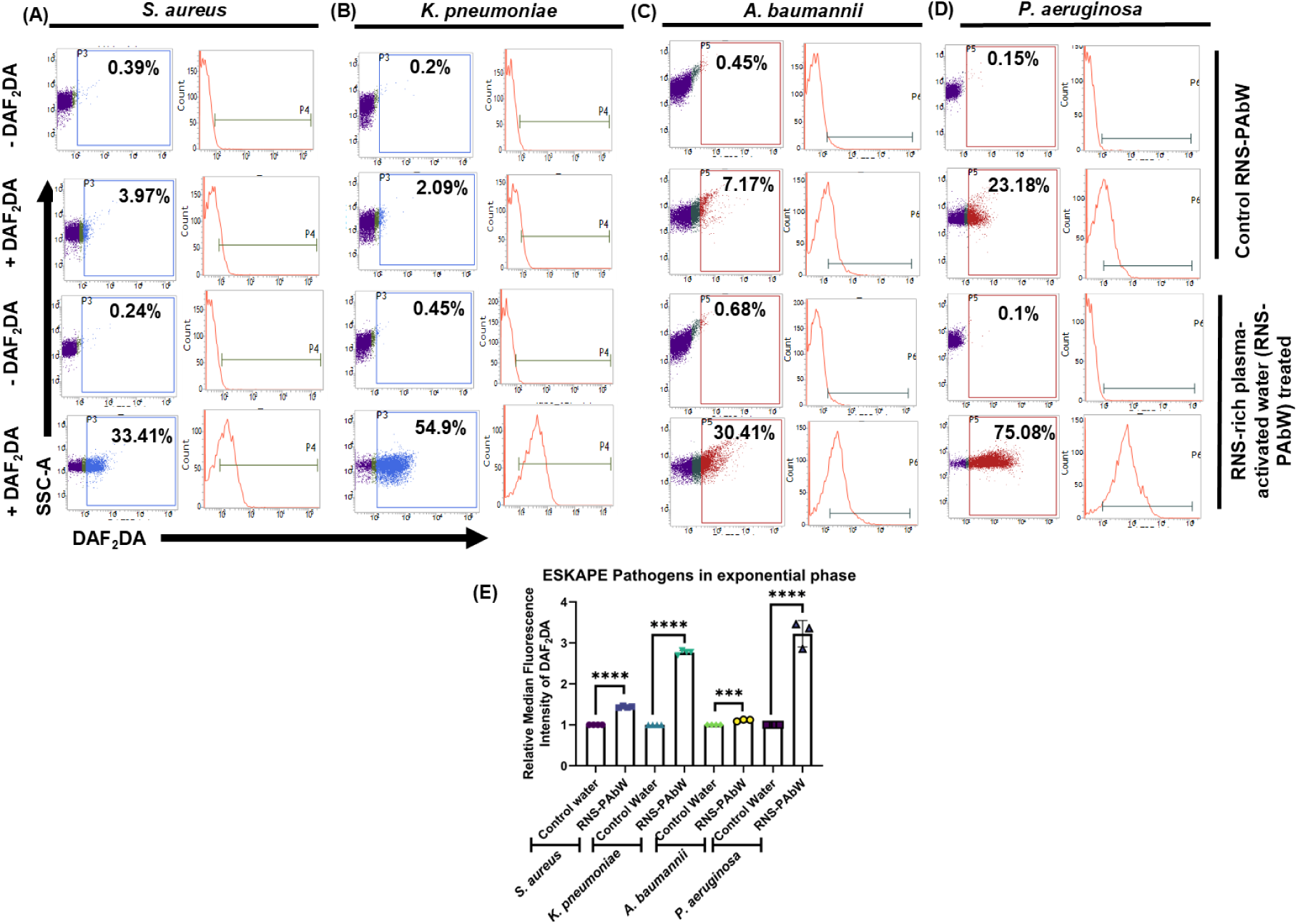
RNS-PAbW generates high amount of reactive nitrogen species in the treated bacteria. (A, B, C, and D) The dot plot (SSC-A vs. DAF_2_DA) and histogram (count vs. DAF_2_DA) represent the RNS content of exponentially growing *S. aureus* (A), *K. pneumoniae* (B), *A. baumannii* (C), and *P. aeruginosa* (D) treated with reactive nitrogen species rich plasma-activated water (RNS-PAbW). The bar graphs represent the median fluorescence intensity (MFI) of the DAF_2_DA-positive population of *S. aureus*, *K. pneumoniae*, *A. baumannii*, and *P. aeruginosa* with or without RNS-PAbW treatment (E). The data are representative of n≥3, N=3 and is presented as mean ± SD. ***(P)* *< 0.05, *(P)* **< 0.005, *(P)* ***< 0.0005, *(P)* ****< 0.0001, ns= non-significant, (Unpaired two-tailed student’s t-test).**

**Figure S4.**
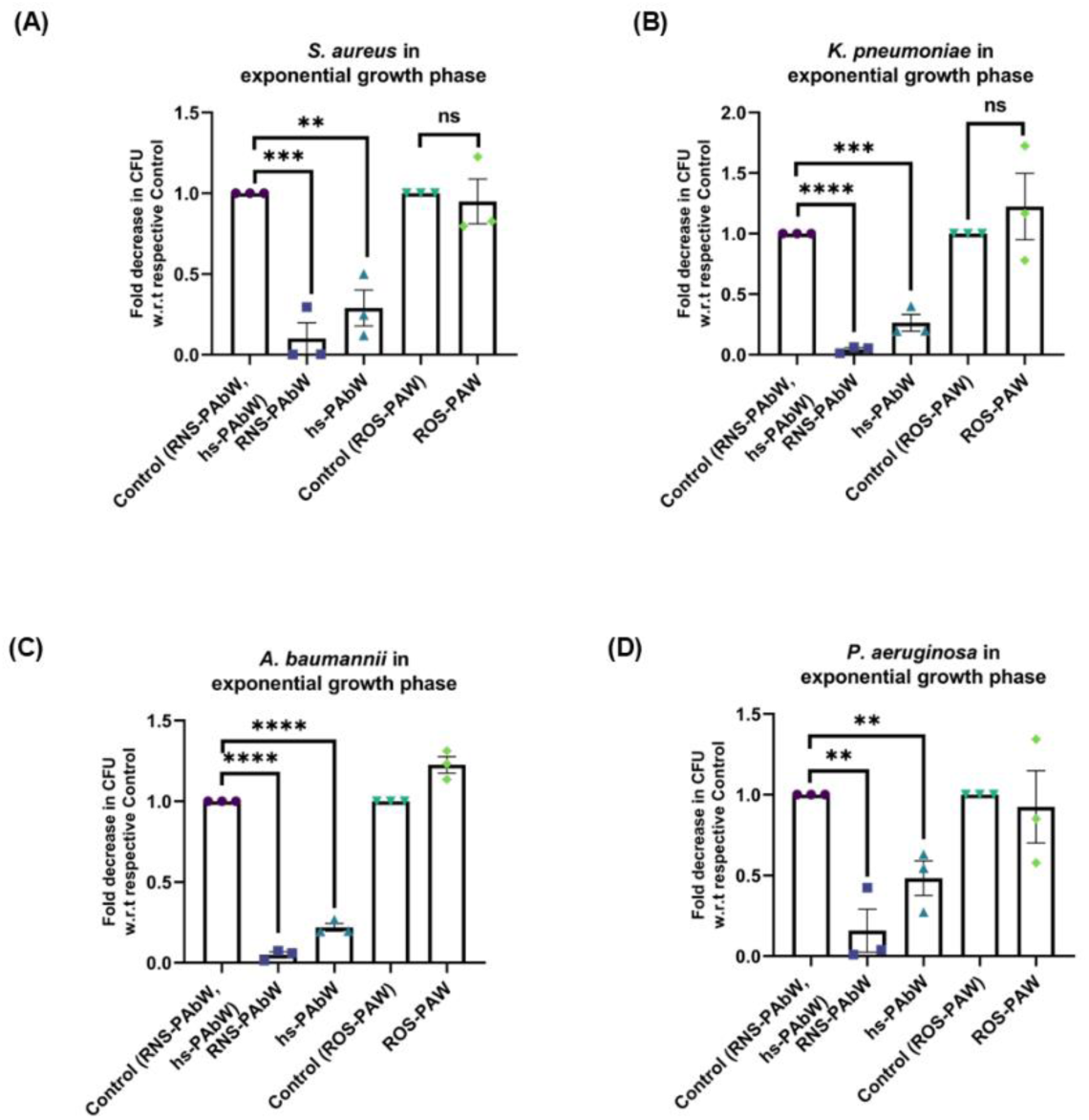
Incubation with RNS-PAbW and hs-PAbW results in significant reduction of Colony Forming Units, but not ROS-PAW. The column graphs represent the fold decrease in CFU after treatment with RNS-PAbW, hs-PAbW, and ROS-PAW compared to the respective control water samples for *S. aureus* (A), *K. pneumoniae* (B), *A. baumannii* (C), and *P. aeruginosa* (D). The data are represented as mean ± SEM. (n=2, N=3). ***(P)* *< 0.05, *(P)* **< 0.005, *(P)* ***< 0.0005, *(P)* ****< 0.0001, ns= non-significant, (Unpaired two-tailed student’s t-test).**

**Figure S5.**
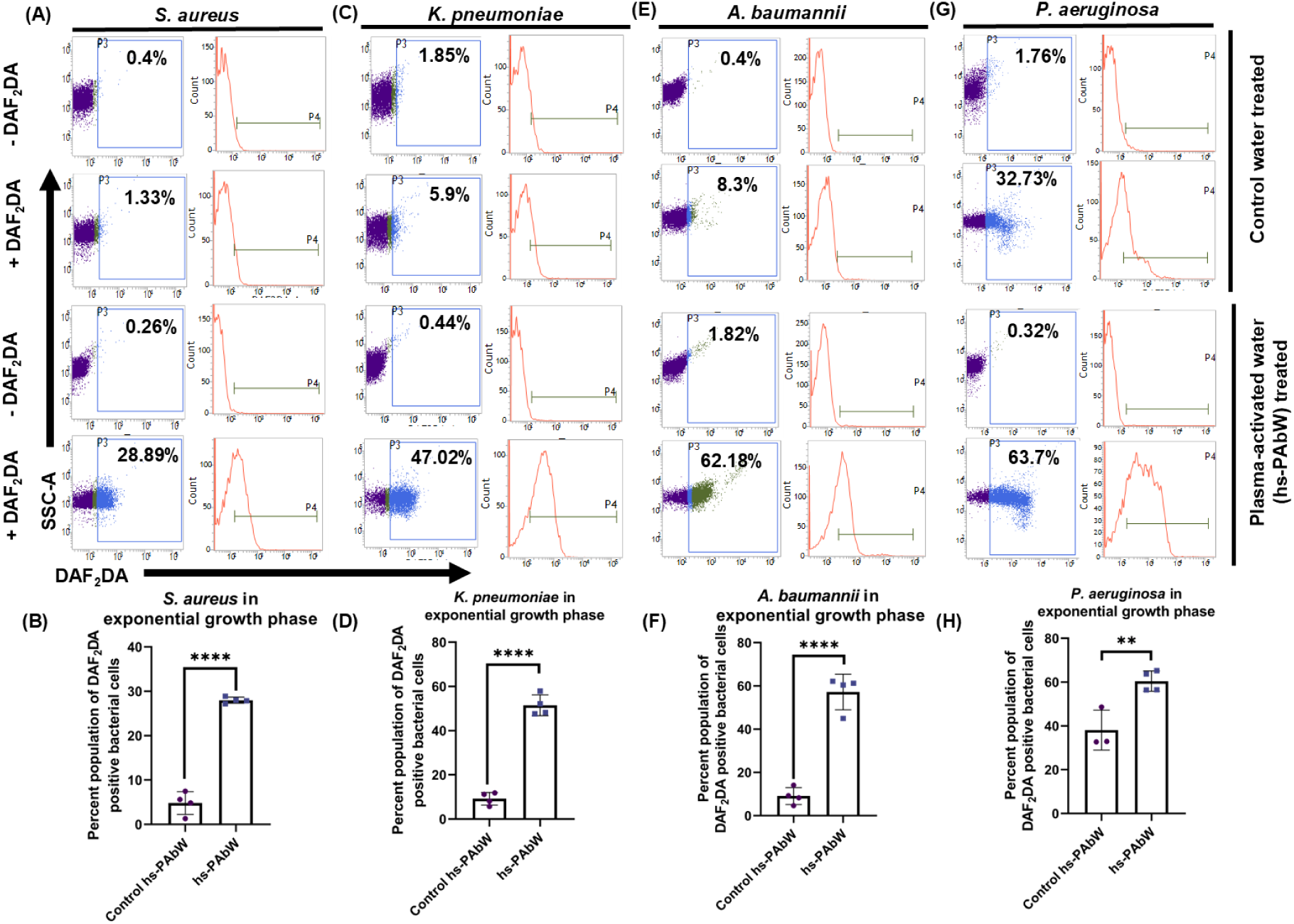
hs-PAbW-dependent generation of reactive nitrogen species while incubated with exponential-phase bacteria. (A, C, E and G) The dot plot (SSC-A vs. DAF_2_DA) and histogram (count vs. DAF_2_DA) represent the generation of reactive nitrogen intermediates upon treating the exponentially growing cultures of *S. aureus* (A), *K. pneumoniae* (C), *A. baumannii* (E), and *P. aeruginosa* (E) with hs-PAbW. The bar graphs represent the DAF_2_DA-positive percent population of *S. aureus* (B), *K. pneumoniae* (D), *A. baumannii* (F), and *P. aeruginosa* (H) with or without hs-PAbW treatment. The data are representative of n≥3, N=3 and is presented as mean ± SD. ***(P)* *< 0.05, *(P)* **< 0.005, *(P)* ***< 0.0005, *(P)* ****< 0.0001, ns= non-significant, (Unpaired two-tailed student’s t-test).**

**Figure S6.**
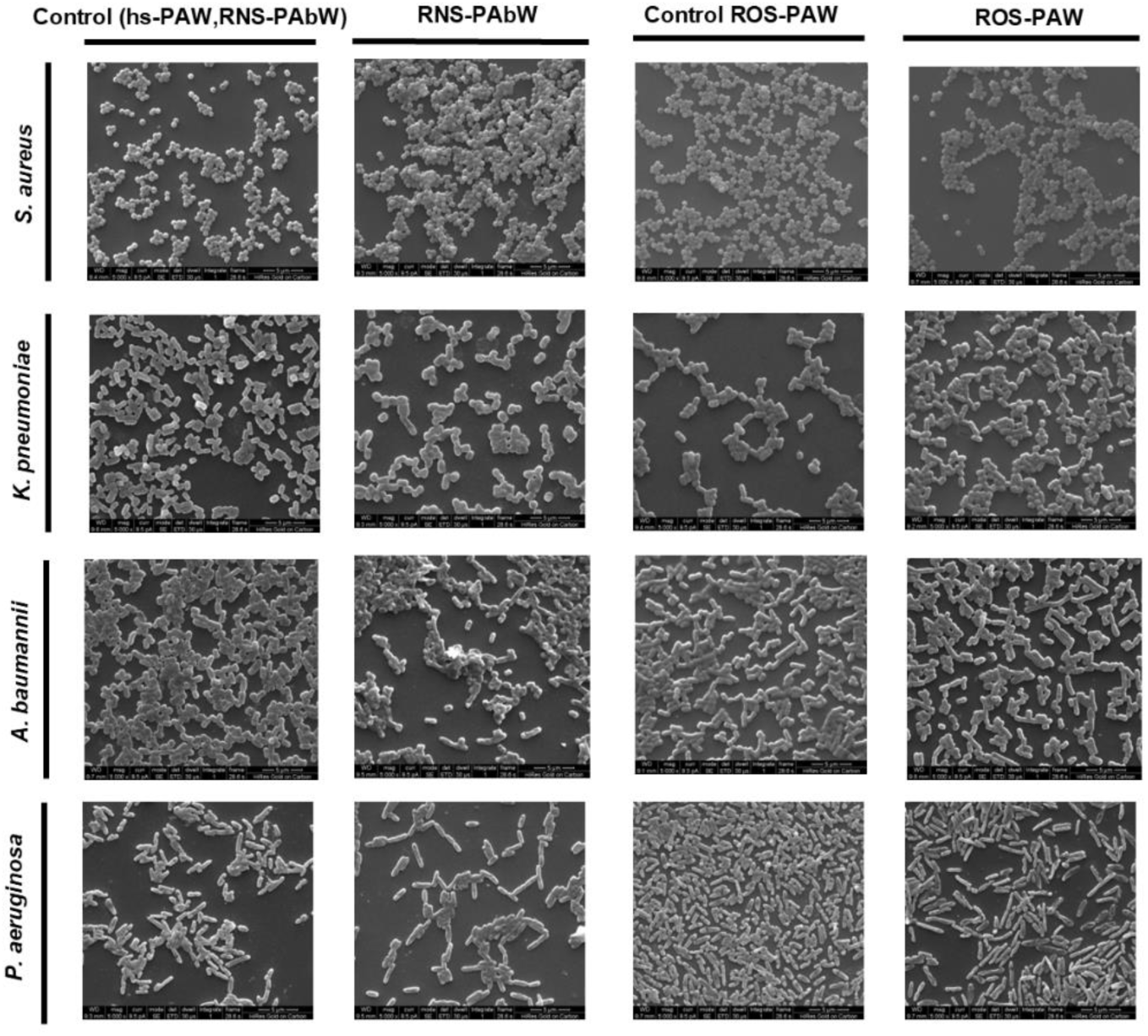
Scanning Electron Microscopy to study morphological change in the bacterial samples post exposure to RNS-PAbW and ROS-PAW. (5000X magnification)

**Figure S7:**
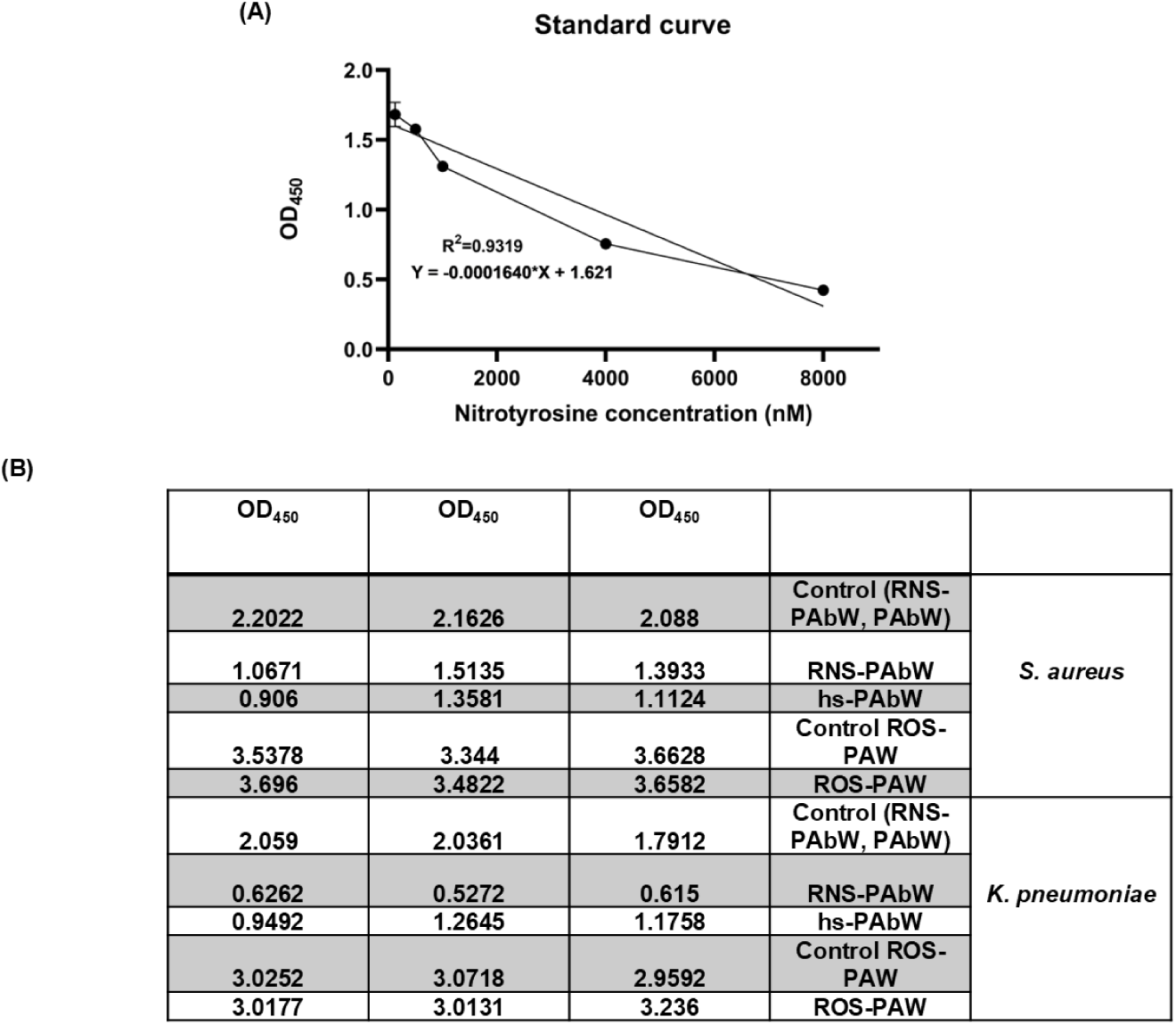
Nitrotyrosine estimation reveals a significant increase in nitrotyrosine levels following treatment with RNS-PAbW and hs-PAbW, but not with ROS-PAW. A standard curve (A) was generated using the OD_450_ absorbance values of the nitrotyrosine standards, and the nitrotyrosine concentration in the samples was determined by interpolating the raw data (B) against the standard curve.

**Figure S8:**
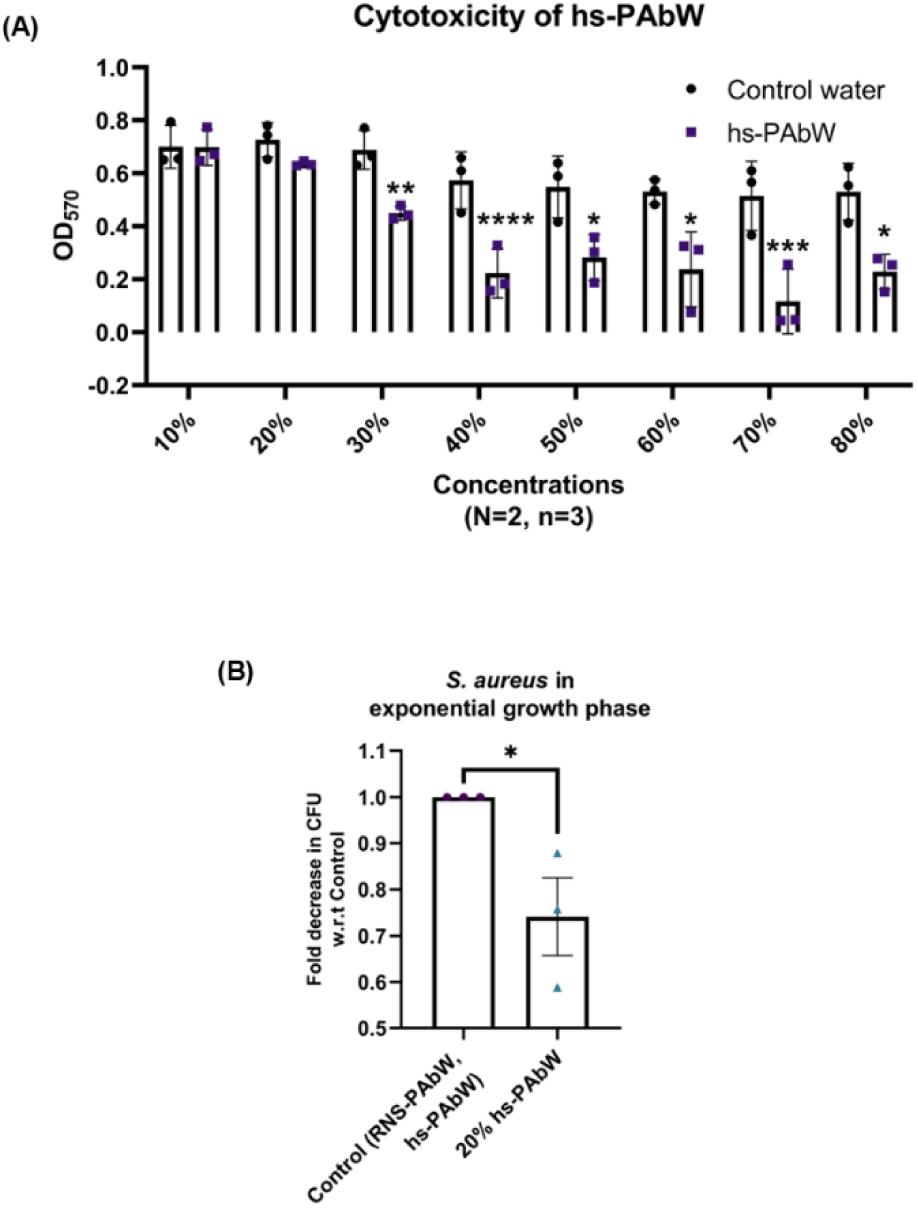
A non-cytotoxic dose of hs-PAbW was also able to retain its bactericidal effect. Figure showing MTT Assay of different concentrations of hs-PAbW (A) and the fold reduction in CFU after treatment with non-cytotoxic dose of hs-PAbW (B). (A) is representative of N=2, n=3 and is expressed as mean± SD and (B) is merged data of N=3, n=2, and is expressed as mean± SEM. ***(P)*\*< 0.05, *(P)* **< 0.005, *(P)* ***< 0.0005, *(P)* ****< 0.0001, ns= non-significant, (One-way ANOVA with mixed effect-analysis for grouped data) (Unpaired two-tailed student’s t-test for column graph).**

